# Bumetanide increases microglia-interneuron contact following traumatic brain injury

**DOI:** 10.1101/2022.06.03.494659

**Authors:** Marine Tessier, Marta Saez Garcia, Emmanuelle Goubert, Li Tian, Florence Molinari, Edith Blasco, Jerome Laurin, François Guillemot, Christian Hübner, Christophe Pellegrino, Claudio Rivera

## Abstract

**Objective:** The Na-K-Cl cotransporter (NKCC1) inhibitor bumetanide has prominent positive effects on the pathophysiology of many neurological disorders. Here we studied whether bumetanide could influence post-traumatic cognitive decline and inflammatory processes by regulating astrocyte and microglia activation.

**Method:** Controlled cortical impacted (CCI) animals were treated with bumetanide during the first post-CCI week. Immunochemistry, flow cytometry, immunoassay, and *in vivo* imaging were used to study astrocytic and microglial morphology and phenotype as well as adult neurogenesis. Telemetric electroencephalograms and cognitive behavioral test were performed at one-month post CCI.

**Results:** Bumetanide prevented CCI-induced decrease in hippocampal neurogenesis and parvalbumin positive interneuron loss. Deletion of NKCC1 in astrocytes neither rescued interneurons nor promote neurogenesis. Interestingly, bumetanide had a strong effect on microglial activation by inducing polarization towards the M1-like phenotype 3 days post-CCI and the M2-like phenotype 7 days post-CCI. Bumetanide increased microglial Brain-derived neurotrophic factor (BDNF) expression and interaction with parvalbumin interneurons. The early treatment with bumetanide resulted in improvements in working and episodic memory, one-month post-CCI, as well as the normalization of theta band oscillations.

**Interpretation:** Here, we disclose a novel mechanism for the neuroprotective action of bumetanide mediated by an acceleration of microglial activation dynamics that leads to an increase of parvalbumin interneuron survival following CCI, possibly resulting from increased microglial BDNF expression and contact with interneurons. Salvage of interneurons may normalize ambient gamma-aminobutyric acid (GABA) resulting in the preservation of adult neurogenesis processes as well as contributing to bumetanide-mediated improvement of cognitive performance.

## Introduction

Traumatic brain injury (TBI) is one of the most prevalent pathologies worldwide with more than 20 million people affected each year ^1^. However, despite being a common disorder, possibilities for intervention to prevent comorbidities and long-lasting sequelae that frequently follow TBI are limited. Understanding the changes and the key players involved in the pathophysiology of the latent phase may help to prevent or reduce some of the comorbidities that accompany TBI.

TBI can frequently be associated with complications such as epileptic seizures, depression and memory impairment ^2^. This may partly result from the alteration of hippocampal information processing and neurogenesis following TBI^3^.^4,5^. However, the mechanisms regulating neurogenesis following TBI remain elusive because of contradictory reports ^4,6^.

Adult neurogenesis is controlled by local GABAergic interneurons and GABA_A_ receptors. In particular, the release of gamma-aminobutyric acid (GABA) by parvalbumin (PV) expressing interneurons controls the quiescent state of radial glia-like cell (RGL) in the dentate gyrus ^7^. Three studies in humans and mouse models of TBI have demonstrated a progressive loss of PV expressing interneurons in both the ipsi- and the contralesional hippocampus, that can persist over months ^8–10^. Goubert et al.^6^ showed a reduction in the number of PV expressing interneurons in injured animals and they demonstrated that the inhibition of chloride uptake by bumetanide, an inhibitor of a Na-K-Cl (NKCC1) co-transport, significantly decreased PV interneuron loss with a positive effect on both depressive-like behavior and cognitive performance. Despite the potential clinical relevance of these results, the underlying mechanisms remain unknown.

Until now, studies on chloride uptake in the brain mainly focused on neuron-intrinsic mechanisms although the expression of the principal cellular uptake transporter NKCC1 is mainly expressed in non-neuronal cells ^11^.

Glial cells are involved early in the temporal sequence of the neuroinflammatory processes following TBI ^12^. Indeed, quiescent glial cells are rapidly activated by a process called “reactive gliosis”. The recruitment of peripheral neutrophils is also observed, which is followed by infiltration of lymphocytes and monocyte-derived macrophages. Simultaneously, the release of pro-inflammatory and anti-inflammatory cytokines promotes and/or inhibits the post-traumatic neuroinflammatory response ^13^. Activated microglia triggers and maintains astrocytic activation through the release of cytokines, which, in turn, act on surrounding glial cells and neurons ^14^. Recent reports showed an important pro-survival function of physical contacts between activated microglia and interneurons, especially PV positive interneurons ^15^.

In this study, we identified a novel mechanism linking cellular chloride uptake in microglia, interneuron survival and neurogenesis that is linked to TBI induced cognitive decline.

## Material and methods

### Animals

The French ethical committee approved all experimental procedures (N°: APAFIS#2797). Wild-type mice had a C57bl6-J background. Three transgenic lines were used : Nestin-GFP ^16^, hGAFP-Cre x NKCC1 flox and B6.129P-Cx3cr1tm1Litt/j ^17^ that were maintained on a mixed genetic background. Only males were used, and they were housed individually in an enriched environment, maintained in a 12 h light/12 h dark cycle environment with controlled temperature (23 ± 2°C), food and water given ad libitum.

### Controlled-cortical impact model (CCI)

Buprenorphine (0.03 mg/kg) was injected intra-peritoneally (i.p) to 10 weeks old C57bl6-J males 30 min before surgery. Mice were then anesthetized using 4 % isoflurane mixed with air and enriched with oxygen (0.3 %) and positioned in a stereotaxic frame (David Kopf Instruments®). Body temperature was monitored throughout the procedure using a rectal probe and maintained at 37 ± 2°C with a heating pad (Harvard Apparatus®). A unilateral craniotomy was performed over the right parietal cortex. CCI was performed using a Leica impactor with standardized parameters (tip diameter 3mm, 6 m/sec speed, 1.5 mm depth and 200 msec duration). Animals were allowed to recover on the heating pad, then were transferred onto post-surgical room. Before the experiment, animals were randomly assigned to one of the following groups: sham-vehicle, sham-bumetanide, CCI-vehicle, and CCI-bumetanide.

### Drug Delivery

Bumetanide stock solution 20 mM (Sigma-aldrich®, B3023) was prepared by dissolving 36.4 mg of powder in 1 ml absolute ethanol. Injections were performed daily twice during the first week post-CCI at 2 mg/kg i.p.

Minocycline (Sigma-Aldrich®, M9511) was injected daily twice, 12 hours apart (45 mg/kg) for one week. The vehicle solution consisted of the same preparation but lacked the bumetanide/Minocycline powder to respect volume and diluent.

### Immunohistochemistry

Mice were transcardially perfused with cold phosphate buffer saline (PBS 1X) prior to 3 % paraformaldehyde solution (AntigenFix, Diapath®). Brains were post-fixed overnight in 3 % paraformaldehyde at 4°C. Sections were permeabilized and blocked in PBS with 0.3 % Triton X-100 and 5 % normal goat serum (NGS) for 1 h at room temperature. Incubation with primary antibodies diluted in PBS with 5 % NGS and 0.3 % Triton X-100 was carried out at 4°C overnight using rabbit anti-DCX (ab18723, Abcam, 1:1000), mouse anti-parvalbumin (p3088, Sigma, 1:500), rabbit anti-Iba1 (W1W019-19741, Wako, 1 :500), mouse anti-GFAP (MAB360, Merck Millipore, 1 :500) and mouse-BDNF antibody (MAB#9, provided by Prof Eero Castren and Yves-Alain Barde). Slices were incubated with the corresponding Alexa Fluor-conjugated secondary antibodies diluted in PBS (Thermo Fisher Scientific, 1 :500) for 2 h at room temperature and finally counterstained for 1 min with Hoechst 33258 (10 μg/mL in PBS, Sigma-Aldrich). Images were taken using a confocal microscope LSM-800 Zeiss (28148). Morphology analysis was performed with the imageJ® plugin Neurphology and contacts quantification with SynapCountJ.

### Flow cytometry analysis

Fresh cortical tissues were dissected and gently chopped and minced in 1mL ice-cold flow cytometry staining buffer (PBS + 1% fetal calf serum) through a 100µm filter on ice. The homogenates were first blocked in PBS with 10% rat serum with gentle rotation in a cold room and then stained for 1h protected from light at 4°C with CD206-FITC (cat no. 141704, Biolegend®) + MHCII-PE (cat no. 107608, Biolegend®) + CD11b-PerCP/Cy5.5 (cat no.101228, Biolegend®) + CD45-APC (cat no.103112, Biolegend®). After staining, cells were fixated with 8% PFA for 30 minutes, centrifuged at 2000rpm. Samples were acquired with a 2-laser, 6 colors cytometer Gallios (Beckman Coulter®). Gating strategy was determined based on the specific staining of markers for microglial population^18^.

### Live Imaging

Five-month-old male mice (B6.129P-Cx3cr1tm1Litt/j, n=5) were anesthetized (ketamine (90mg/kg) and xylazine (4.5 mg/kg), i.p). Animals received subcutaneous injections of dexamethasone sodium phosphate (2ug/g) and carprofen (5mg/ml). The bone fragment was removed using a 0.45 mm diameter drill. A coverslip was placed on top of the cranial window. A metallic head-plate was attached to the coverslip forming a frame for the window. Animals were monitored during the following week to make sure the recovery was successful.

One week before imaging, animals were trained in the Mobile HomeCage® (MHC V5, Neurotar®). Training lasted 5 days.

One month after surgery, trained animals were placed under the 2-photon microscope (Femtonics®) and imaged for 4 days (Injury Day, 1 day post injury, 3 days post injury and 5 days post injury).

Each animal had first a 30 min baseline recording of 100 µm. After the injury, z-stacks covered the entire injury area (around 300 µm). Z-stacks were collected every 5min for 1h. Injury was achieved by exposing 1 cell for 20s at 75 us/pixel and 100% laser power.

To evaluate the surveillance state of microglia *in vivo* we measured the processes length of microglial cells and its evolution at 1- and 5-days post injury. We also estimated the dendritic surveillance (motility) by quantifying the standard deviation of the processes’ changes in length during the recording time window.

### Cell culture

As described previously ^19^, BV2 cells were cultured in DMEM media containing 10 % fetal bovine serum, 100 IU/mL penicillin, and 100 mg/mL streptomycin for amplification. After seeding, we cultured BV2 cells in DMEM media containing only 1% fetal bovine serum, 100 IU/mL penicillin, and 100 mg/mL streptomycin. Serum deprivation allows cells differentiation. Plates were incubated at 5% CO2, 37 °C. LPS (50 ng/mL), bumetanide (40 nM) or LPS + Bumetanide were applied in the culture media for either 24 or 72 hours.

### Western blotting

Western blot was performed as described in Goubert et al. ^6^ Membranes were exposed overnight at 4°C to NKCC1-specific antibody (DSHB Hybridoma Product T4) diluted 1:2000 in blocking solution. Chemiluminescent detection was performed using ECL-plus kit (Pierce Biotech®). We measured signal intensities with the image analysis software G box (Syngene®). Then, membranes were stripped, and probed with rabbit anti-α-tubulin (Sigma®, 1:10000). Quantification was performed using Gel Plot Analyser plugin (Fiji®).

### ELISA

Quantification of mature BDNF (mBDNF) was performed with mBDNF Rapid ELISA Kit (Biosensis®, BEK-2211-1P/2P – sandwich ELISA -Thebarton, SA, Australia) following the manufacturer’s protocol. Quality Control sample ranged between 175-325 pg.ml-1 and the mBDNF standard ranged between 7.8-500 pg.ml-1. Concentrations were determined using the FLUOstar® OPTIMA microplate reader (BMG Labtech, France). Microglia interleukines production was analyzed the same way with the Mouse Interleukin 6 ELISA Kit (Biosensis®, BEK-2043-1P) and the Mouse Interleukin 10 ELISA Kit (Biosensis®, BEK-2046-1P)

### Behavioral tests

One-month post-CCI, mice were tested on 2 different tasks using the object recognition paradigm to test the individual components of episodic-like memory. The novel object recognition (NOR) and the object displacement task (ODT) ^20^. Animals were habituated to the testing room 24 h before testing. For the NOR, mice were placed in the center of maze (Noldus apparatus®, 38.5 cm x 38.5 cm) and allowed to freely explore the space for 10 min, then two identical objects were added for a 3-min exploration time, finally after a 3-min retention time, one of the objects was replaced and the time of exploration was measured for a 3-min period. This test was used to assess the short-term memory. The same experiment was also performed with 1-hour retention time to test the long-term memory. For the ODT, mice were placed in the center of the same maze, for 3 min, then two identical objects were added for 3 min. After 3min retention time, one of the objects was moved and the time of exploration of each object was recorded. Recording and analyses were done using the Ethovision software (Noldus®).

### In vivo electrophysiological recordings

Scalp electroencephalographic (EEG) recordings were performed in freely moving mice 3 weeks post-CCI. Telemetric recording electrodes were implanted 2mm posterior to and 1.5mm left of the bregma, in the parietal cortex. A reference electrode was placed rostral in the cerebellum. After a 72hour recovery, EEG (amplified 31,000, filtered at 0.1–120Hz pass, acquired at 1000Hz) was monitored using a telemetric system (EMKA TECHNOLOGIES S.A.S) for 3 days, 24 hours per day. Recordings were analyzed offline using the “Power spectra for continuous” module of NeuroExplorer version 5.305.

## Results

### Bumetanide treatment rescues memory impairment and theta band oscillations

Memory impairment following TBI is characterized by changes in theta rhythms ^21,22^. Using telemetric EEG, we found that at one-month post-CCI the power of theta band oscillations in vehicle treated animals is significantly lower compared to sham (Sham 30.44 % ± 0.36 vs CCI Veh 21.14 % ± 4,82, Fig 1B, C and D). We also found that bumetanide treatment normalized the contribution of theta band oscillations to the power spectra and was significantly different from vehicle treated animals (CCI Bum 29.10 % ± 4.41, Fig 1B, C and D).

**FIG 1.**
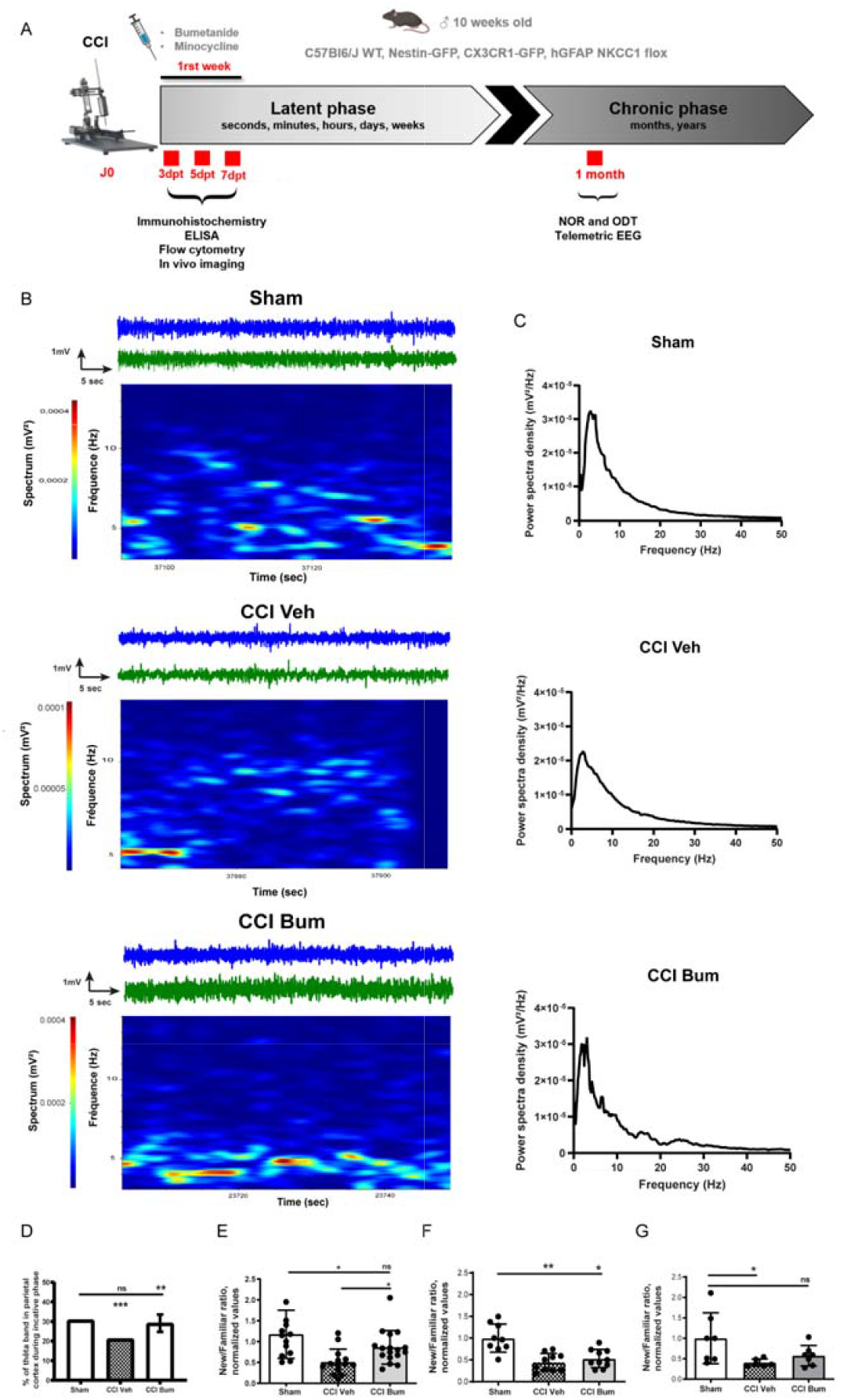
Bumetanide treatment improves CCI induced behavioral changes. **(A)** Schematic representation of the experimental timeline. **(B)** Morlet wavelet transform of example recording session from a Sham, CCI Veh and CCI bum animals during inactive period. Blue traces are raw data, and green traces are band passed for theta band (3 to 7 Hz). **(C)** Example of EEG power spectral density for each animal, n = 5, 6 and 6 respectively. **(D)** The percentage of theta band in the contralesional parietal cortex of mice 1 month post TBI is modified, bumetanide treatment rescues it. **(E)** The Novel Object Recognition task after 1hour retention time. **(F)** The Novel Object Recognition task after 3min retention time. **(G)** The Object Displacement Task. For all experiments, the results are presented on a ratio of time of new versus familiar, n = 10, 11 and 10 respectively.

Post TBI changes in EEG are correlated with behavioral changes^23^. To investigate a possible effect of bumetanide on post-TBI cognitive behavior we performed two cognitive performance test, Novel Object Recognition (NOR) and Object Displacement Task (ODT). CCI induced significant decrease in the recognition ratio in all task examined (ODT normalized to Sham: CCI Veh 0.38 ± 0.06 ; NOR 5 min normalized to Sham CCI Veh 0.44 ± 0.20; NOR 1h Sham 1.17 ± 0.58 vs CCI Veh 0.507 ± 0.31, Fig 1E, F and G). Interestingly the strongest effect of bumetanide was found in NOR task using 1h retention (CCI Bum 0.86 ± 0.40, Fig 1E).

### Bumetanide rescues CCI-induced changes in secondary neurogenesis and PV interneuron loss

Interneurons are important contributors to pathological brain oscillations^24^. Bumetanide-induced changes in Theta power could reflect partially changes in interneuron subpopulations. Parvalbumin positive interneurons are a particularly vulnerable subpopulation following CCI^7^. We quantified PV-positive interneurons in the granular layer of the ipsi and contralesional DG during the first post-traumatic week at 3 and 7 d after CCI. Both sides showed a significant reduction of the number of PV interneurons at 3 days after CCI (normalized value on sham: sham contra 1.00 ± 0.20 vs contra Veh 0.52 ± 0.18 and sham ipsi 1.00 ± 0.22 vs ipsi Veh 0.22 ± 0.18, Figs 2A and B). This loss was significantly reduced by bumetanide treatment in the contra-(0.77 ± 0.23) and ipsilesional sides (0.51 ± 0.14, Fig 2A and B). We also quantified the PV-containing interneurons survival at 7 d after CCI and also found a significant reduction in both sides (normalized value on sham: sham contra 1.0 ± 0.27 vs contra Veh 0.59 ± 0.18 and sham ipsi 1.00 ± 0.27 vs ipsi Veh 0.14 ± 0.12, Fig 2C and D). This loss was significantly reduced by bumetanide treatment at the contra-(1.02 ± 0.34) and ipsilesional side (0.71 ± 0.24, Fig 2C and D).

**FIG 2.**
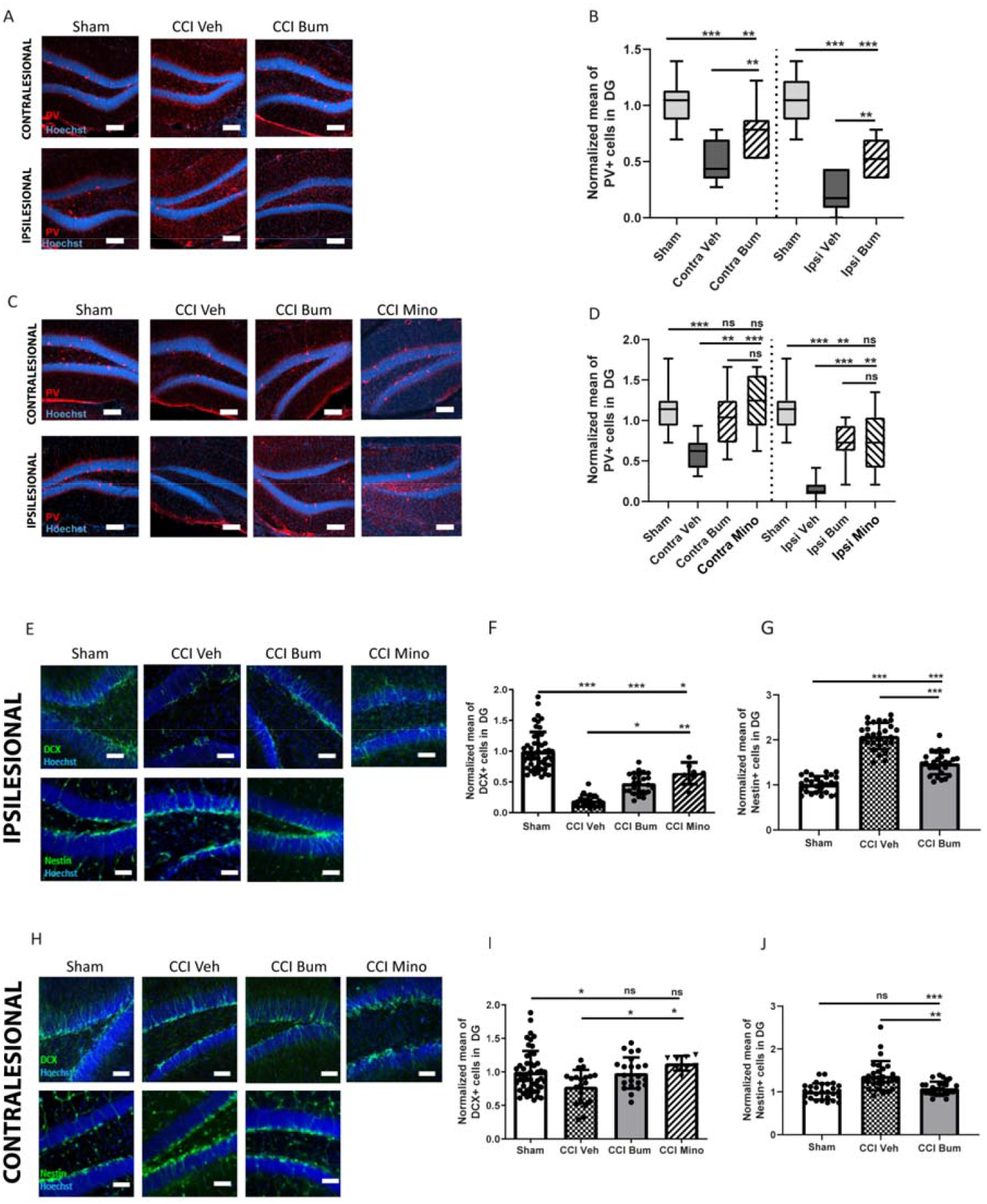
Effect of bumetanide and minocycline on CCI-induced changes in adult neurogenesis and PV interneuronal loss in the dentate gyrus. **(A)** Parvalbumin (PV) labeling at 3 days post-CCI in the contralesional (upper pictures) and ipsilesional (lower pictures) dentate gyrus of sham, CCI vehicle and CCI bumetanide-treated animals (scale bar = 100 µm). **(B)** Quantification of PV positive cells 3dpCCI in the contralesional and ipsilesional dentate gyrus of sham, CCI vehicle and CCI bumetanide-treated animals. **(C)** Same as **(A)** at 7 dpCCI with minocycline-treated animals added. **(D)** Same as **(B)** 7dpCCI. **(E)** Doublecortin (DCX, upper pictures) and Nestin (lower pictures) labeling at 7 days post-CCI in the contralesional dentate gyrus of sham, CCI vehicle, bumetanide-treated and minocycline-treated animals (scale bar = 100 µm). **(F)** Quantification of DCX positive cells 7dpCCI in the contralesional dentate gyrus of sham, CCI vehicle, bumetanide-treated and minocycline-treated animals. **(G)** Quantification of Nestin positive cells 7dpCCI in the contralesional dentate gyrus of sham, CCI vehicle, bumetanide-treated and minocycline-treated animals. **(H)** Same as in **(E)** in the ipsilesional dentate gyrus. **(I)** Same as in **(F)** in the ipsilesional dentate gyrus. **(J)** Same as in **(G)** in the ipsilesional dentate gyrus.DCX, Nestin and PV stainings : n = 7 animals per condition, 3 slices per animal. All sets of data were analyzed using Brown-Forsythe Anova test with Dunnett’s post hoc test. *p < 0.05; **p < 0.01; ***p < 0.001.

Ambient GABA, provided by the ongoing activity of DG interneurons and particularly PV interneurons, plays a role in the proliferation and migration of granular cell progenitors ^6,25^. A loss of PV interneurons may significantly contribute to the TBI-induced changes in stem cell proliferation by reducing GABA release. Seven days post-CCI, we observed a significant reduction in the number of DCX positive neurons within both, the ipsi- and contralesional DG (normalized value on sham: sham ipsi 1.00 ± 0.31 vs. CCI Veh ipsi 0.19 ± 0.09 and sham contra 1.00 ± 0.31 vs. CCI Veh contra 0.78 ± 0.24, Figs 2E, F, H and I). This coincided with an increase in the number of Nestin positive cells within the DG (sham ipsi 1.00 ± 0.17 vs. CCI Veh ipsi 2.08 ± 0.29 and sham contra 1.00 ± 0.18 vs. CCI Veh contra 1.37 ± 0.34 Figs 2E, G, H and J). Bumetanide treatment reduced the number of Nestin positive cells (CCI Bum ipsi 1.4 ± 0.24 and CCI Bum contra 1.0 ± 0.15, Figs 2E, G, H and J) and triggered an increase in the number of DCX-positive cells (CCI Bum ipsi 0.47 ± 0.16 and CCI Bum contra 0.98 ± 0.23, Figs 2E, F, H and I).

TBI is known to induce strong inflammatory reaction. Interestingly, treatment with the anti-inflammatory agent minocycline resulted in identical effects on both PV-survival (Contra Veh 0.59 ± 0.18 vs Contra Mino 1.21 ± 0.34 and ipsilesional side Ipsi Veh 0.14 ± 0.12 vs Ipsi Mino 0.74 ± 0.39) and newborn neuron (CCI Mino ipsi 0.63 ± 0.17 and CCI Mino contra 1.12 ± 010, Figs 2A, B, C and D). These results suggest that part of the positive effects of bumetanide on interneuron survival and proliferation are mediated trough an anti-inflammatory mechanism.

### The positive effects of bumetanide on CCI outcome are not mediated via astrocytes

Neuroinflammation is present in the primary (acute) and secondary (chronic) stages of TBI ^12^ where microglia and astrocytes activation are involved in the mechanisms leading to both adverse and beneficial effects. Under physiological conditions NKCC1 is predominantly expressed in astrocytes and microglia compared to neurons ^26^. Therefore, we wondered whether the effects of bumetanide are mediated by these cell populations. We did not observe any effect of bumetanide on astrocytes morphology on both sides at 3 and 5 days after treatment (data not shown). We only found significant effects on the contralateral side at 7 days post CCI where Bumetanide significantly decreased the length of astrocyte processes 7 days post-CCI in the contralesional DG (CCI Veh 290 µm ± 10.91 vs CCI Bum 147 µm ± 22.17, Fig 3A). The treatment also decreased the soma’s area (Sham 1951 µm^2^ ± 633.6 vs CCI Veh 3232 µm^2^ ± 805 vs CCI Bum 1888 µm^2^ ± 465.8) and the number of endpoints (Sham 895.5 ± 108,2 vs CCI Veh 594.4 ± 130.4 vs CCI Bum 957.2 ± 155.9; Fig 3A and B).

**FIG 3.**
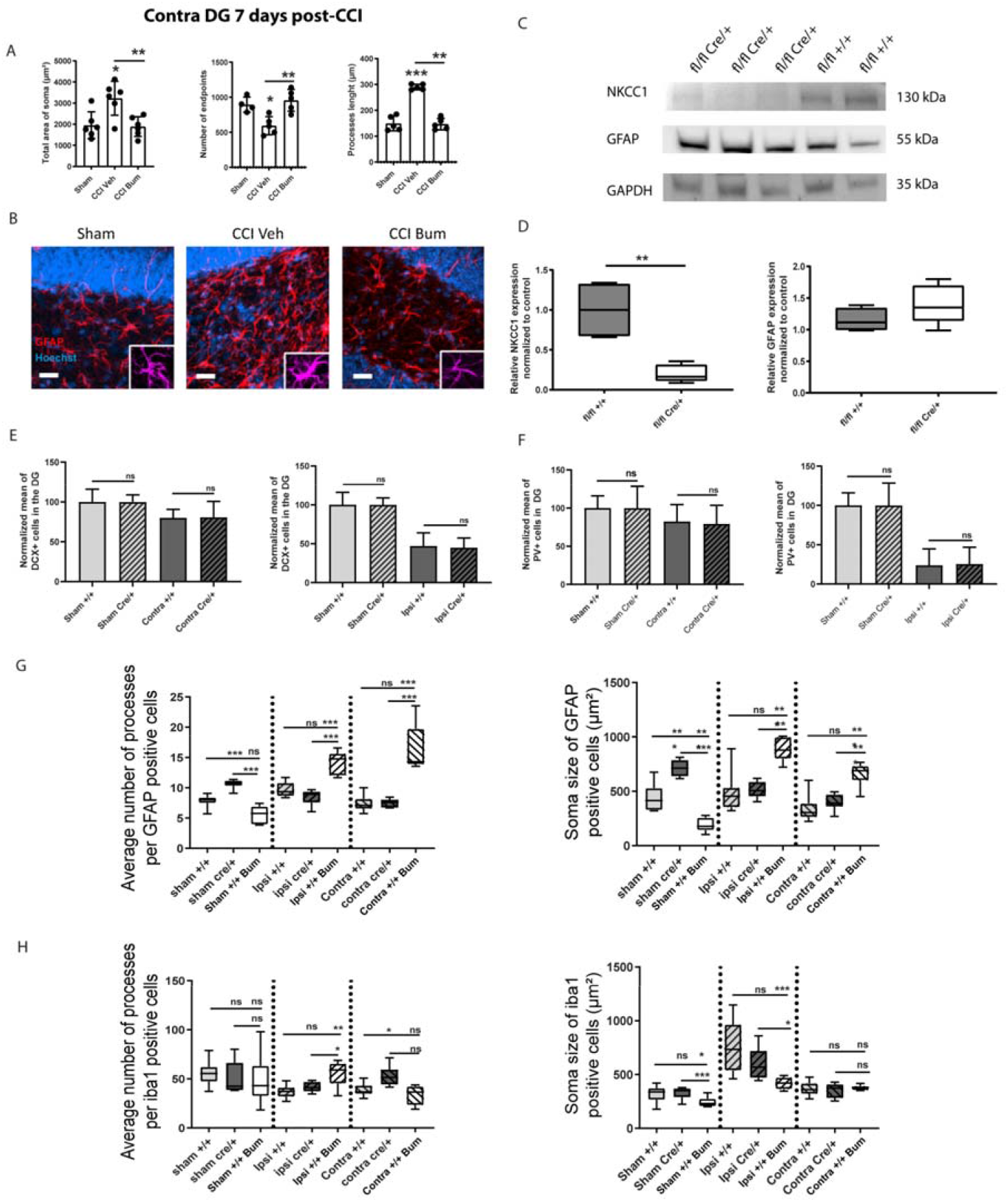
Astrocytes morphological changes after CCI in the contralesional side 7 days post-CCI and effect of bumetanide on main inflammation cells. **A)** Total area of GFAP+ cells soma, quantifications of GFAP processes endpoints’, attachment points and lenght in the contralesional dentate gyrus, 7 days post-CCI. n=5 animals, 2 slices per animal. **(B)** GFAP immunostaining from sham, CCI-vehicle and CCI-bumetanide treated animals in the contralesional dentate gyrus 7 days post-CCI. **(C)** and **(D)** NKCC1 and GFAP protein expression normalized to the ubiquitous marker GAPDH. **(E)** Quantification of DCX positive cells 7dpCCI in the dentate gyrus of sham NKCC1 fl/fl GFAP +/+, sham NKCC1 fl/fl GFAP Cre/+, CCI NKCC1 fl/fl GFAP +/+ and CCI NKCC1 fl/fl GFAP Cre/+. **(F)** Quantification of PV positive cells 7dpCCI in the dentate gyrus of sham NKCC1 fl/fl GFAP +/+, sham NKCC1 fl/fl GFAP Cre/+, CCI NKCC1 fl/fl GFAP +/+ and CCI NKCC1 fl/fl GFAP Cre/+. **(G)** Quantification of number of processes and average soma’ size of astrocytes (GFAP+ cells) in NKCC1 fl/fl GFAP-Cre/+ and NKCC1 fl/fl GFAP +/+ mice at 7 days post-CCI. **(H)** Quantification of number of processes and average soma’ size of microglia (Iba1+ cells) in NKCC1 fl/fl GFAP-Cre/+ and NKCC1 fl/fl GFAP +/+ mice at 7 days post-CCI. Morphological analyses were quantified using the ImageJ plugin Neurphology. All sets of data were analyzed using Brown-Forsythe Anova test with Dunnett’s post hoc test or using a t-test with Welch’s correction. *p < 0.05; **p < 0.01; ***p < 0.001.

To investigate whether the effect of bumetanide on adult neurogenesis depends on NKCC1 in astrocytes, we monitored the effect of CCI in transgenic mice in which NKCC1 was specifically deleted in astrocytes (GFAP-NKCC1 KO), thus mimicking the effect of bumetanide on astrocytes (Fig 2C and D). Interestingly, there was no impact on adult neurogenesis in GFAP-NKCC1 KO mice, neither in the contralesional DG (Sham +/+ 100 % ± 16.03 vs Sham Cre/+ 100 % ± 8.91 and Contra +/+ 80.1 % ± 10.6 vs Contra Cre/+ 80.7 % ± 19.90) nor in the ipsilesional DG (Ipsi +/+ 47.1 % ± 16.85 vs Ipsi Cre/+ 44.9 % ± 12.46, Fig 3E). Similarly, we found no effect of NKCC1 deletion in astrocytes in CCI-induced PV interneuron decrease in both the contra-(Sham +/+ 100 % ± 15,96 vs Sham Cre/+ 100,00 % ± 28,43 and Contra +/+ 82,34 % ± 22,20 vs Contra Cre/+ 79,34 % ± 24,22, Fig 3F) and ipsilesional DG (Ipsi +/+ 23,94 % ± 20,81 vs Ipsi Cre/+ 25,44 % ± 21,40), (Fig 3F).

We also compared the effect of bumetanide treatment and NKCC1 deletion in astrocytes in GFAP-NKCC1 knock-out (KO) mice 7 days post-CCI. While bumetanide had no effect on the number of processes in sham WT animals (Sham +/+ 7.90 ± 0.77 vs Sham +/+ Bum 5.55 ± 1.51) it caused a significant increase in both the ipsi- and the contralesional DG (Ipsi +/+ 9.76 ± 1.21 vs Ipsi +/+ Bum 14.10 ± 1.98 and Contra +/+ 7.37 ± 1.16 vs Contra +/+ Bum 16.76 ± 4.02), GFAP-KCC1 KO animals showed an increase of the number of processes in sham (Sham +/+ 7.90 ± 0.77 vs Sham Cre/+ 10.64 ± 0.73) and no effect in both the ipsi- and contralesional side after Bumetanide treatment (Ipsi +/+ 9,76 ± 1,21 vs Ipsi Cre/+ 8,45 ± 1,22 and Contra +/+ 7.37 ± 1.16 vs Contra Cre/+ 7.52 ±0.68), (Fig 3G). In addition, bumetanide treatment induced a decrease of astrocyte soma size in WT controls (Sham +/+ 438.2 µm^2^ ± 121.4 vs Sham +/+ Bum 193.1 µm^2^ ± 67.36) and a significant increase in both the ipsi- and contralesional side after CCI (Ipsi +/+ 495.4 µm^2^ ± 189.7 vs Ipsi +/+ Bum 885.7 µm^2^ ± 112.5 and Contra +/+ 331.6 µm^2^ ± 103.6 vs Contra +/+ Bum 655.9 µm^2^ ± 114.9). Conversely, in GFAP-NKCC1 KO we observed a significant increase of astrocyte soma size in sham animals (Sham +/+ 438.2 µm^2^ ± 121.4 vs Sham Cre/+ 713.3 µm^2^ ± 73.82) and no changes neither in the ipsi-nor contralesional side after CCI (Ipsi +/+ 495.4 µm^2^ ± 189.7 vs Ipsi Cre/+ 514.0 µm^2^ ± 73.69 and Contra +/+ 331.6 µm^2^ ± 103.6 vs Contra Cre/+ 395.2 µm^2^ ± 73.48), (Fig 3G).

There is a complex interplay between astrocyte and microglia activation during pathophysiological conditions. For this reason, we also investigated the effect of GFAP-NKCC1 KO on microglia. At 7 days of bumetanide treatment the number of processes in microglia in WT animals did not differ either between sham nor contralesional DG of CCI animals (Sham +/+ 55.53 µm ± 12.56 vs Sham +/+ Bum 49.11 µm ± 24,52 and Contra +/+ 38.82 µm ± 6.241 vs Contra +/+ Bum 34.01µm ± 9.768), (Fig. 3H). However, it led to an increase of the number of microglia processes in the ipsilesional side (Ipsi +/+ 36.50 ± 6.025 vs Ipsi +/+ Bum 55.57 ± 12.40). In GFAP-NKCC1 KO mice, there was no change in the number of processes in both sham condition (Sham +/+ 55.53 ± 12.56 vs Sham Cre/+ 50.92 ± 16.25) and in the ipsilesional side (Ipsi +/+ 36.50 ± 6.025 vs Ipsi Cre/+ 41.65 ± 4.984) of CCI animals. Though, a significant increase was observed in the contralesional side of CCI animals (Contra +/+ 38.82 ± 6.241 vs Contra Cre/+ 52.78 ± 10.42). The average size of microglia’s soma was significantly reduced after 7 days of bumetanide treatment in sham WT mice (Sham +/+ 318,1 µm^2^ ± 70.85 vs Sham +/+ Bum 245.5 µm^2^ ± 47.64) in the ipsilesional side (Ipsi +/+ 750.3 µm^2^ ± 227.0 vs Ipsi +/+ Bum 419,6 µm^2^ ± 54.48) but there were no changes in the contralesional side (Contra +/+ 367.7 µm^2^ ± 65.99 vs Contra +/+ Bum 379.0 µm^2^ ± 23.37). In GFAP-NKCC1 KO animals there were no changes neither in sham condition (Sham +/+ 318.1 µm^2^ ± 70.85 vs Sham Cre/+ 329.2 µm^2^ ± 53.61) nor in the ipsi-(Ipsi +/+ 750.3 µm^2^ ± 227.0 vs Ipsi Cre/+ 597.0 µm^2^ ± 145.7) or contralesional side (Contra +/+ 367.7 µm^2^ ± 65.99 vs Contra Cre/+ 353.8 µm^2^ ± 68.58), (Fig 3H).

These results indicated that the depletion of NKCC1 in astrocytes leads significantly different effects on astrocyte morphology than bumetanide treatment. In addition, the positive effects that occur in neurogenesis and interneuron survival after bumetanide treatment might not be mediated by astrocytes.

### Bumetanide rescues the CCI induced loss of PV interneurons by changing microglia morphology and phenotype

We then wondered if bumetanide could influence microglia morphology at early stages post-CCI. Bumetanide treatment induced a significant increase of microglia endpoint numbers (Sham 715.9 ± 81.03 vs CCI Veh 698.8 ± 331.2 vs CCI Bum 1076.0 ± 85.13), number of attachment points (Sham 41.86 ± 12.40 vs CCI Veh 101.9 ± 43.70 vs CCI Bum 140.8 ± 54.96) and soma’s area (Sham 1458 µm^2^ ± 652.7 vs CCI Veh 3494 µm^2^ ± 757.6 vs CCI Bum 4985 µm^2^ ± 850,6) (Fig 4A and B) in the ipsilesional DG 3 days post-CCI.

**FIG 4.**
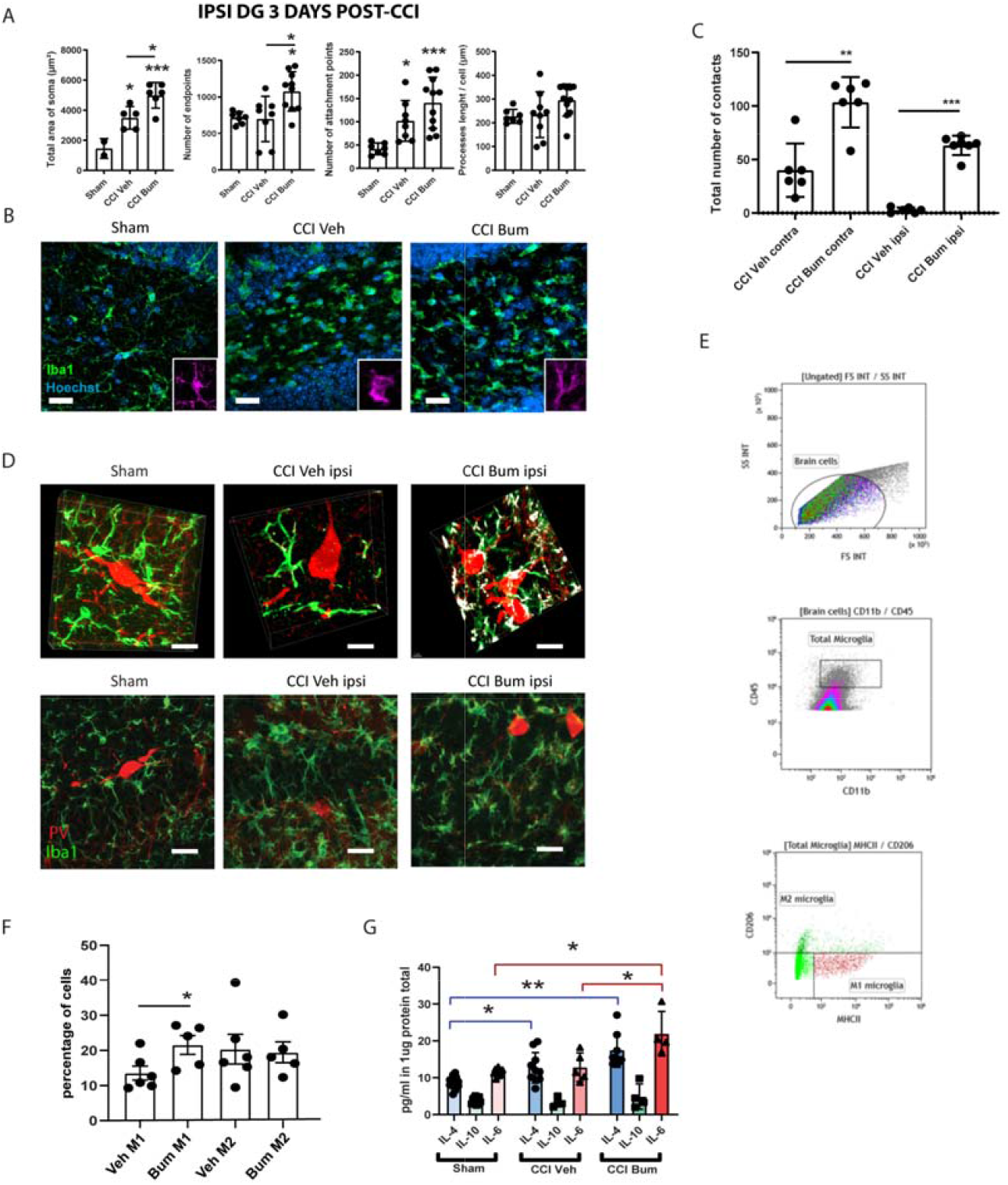
Microglia phenotypes and morphological changes after CCI and effect of bumetanide in the ipsilesional dentate gyrus at 3 days post-CCI. **(A**) Total area of Iba1+ cells soma, quantifications of Iba1 processes endpoints’, attachment points and length in the ipsilesional dentate gyrus. n=5 animals, 2 slices per animal. **(B)** Iba1 immunostaining from sham, CCI-vehicle and CCI-bumetanide treated animals. Purple pictures were extracted from different pictures. **(C)** Quantification of the number of contacts between microglia (Iba1 staining) and parvalbumin interneurons at 3 days post-CCI. n=5 animals per condition, 3 slices per animal. **(D)** Example of parvalbumin and Iba1 immunostaining, from sham, CCI-vehicle and CCI-bumetanide treated animals, in the ipsilesional dentate gyrus, 3D representations (upper pictures), and stack (lower pictures). **(E)** Flow cytometry gating plots at 3 days post-CCI in ipsilesional side. **(F)** Analysis of microglia phenotypes by calculating the percentage of CD45 ^low^ CD11b^+^ MHCII^high^ CD206^low^ (M1) and CD45^low^ CD11b^+^ CD206^high^ MHCII^int^ (M2) markers by flow cytometry at 3 days post-CCI. n=5-6 animals per condition. **(G)** Quantity of IL-4, IL-10 and IL-6 produced in the ipsilesional hippocampus. Phenotype changes were analysed by Welch Anova followed by T-test. Morphological analysis was quantified using the ImageJ plugins Neurphology and SynapCountJ and analyzed using a t-test with Welch’s correction. Interleukin production was analyzed using a Kruskall-Wallis test for IL-10 *p < 0.05; **p < 0.01; ***p < 0.001.

A recent study showed that microglia process motility and their interaction with interneurons play an important role in PV survival ^15^. Hence, we quantified the number of contacts between microglial cells and PV interneurons. We found significant differences between vehicle and bumetanide-treated animals at 3 days post CCI in both the contra-(CCI Veh 40.00 ± 24.92 vs CCI Bum 103.50 ± 23.55) and ipsilesional side (CCI Veh 2.75 ± 2.48 vs CCI Bum 63.20 ± 9.04) (Fig 4C and D).

Using flow cytometry, we found that microglia were more likely to acquire the pro-inflammatory phenotype M1 (Veh M1 13.55 % ± 4.91 vs Bum M1 21.59 % ± 6.03), (Fig 4E and F) after 3 days of bumetanide treatment. To further confirm this, we measured interleukins in brain tissue and found a significant increase in IL-6 in the hippocampus of CCI animals treated with bumetanide (CCI Bum 21.94 pg/mL ± 6.03 vs Sham 11.49 pg/mL ± 0.91 and CCI Veh 12.85 pg/mL ± 3.89, Fig 5G).

**FIG 5.**
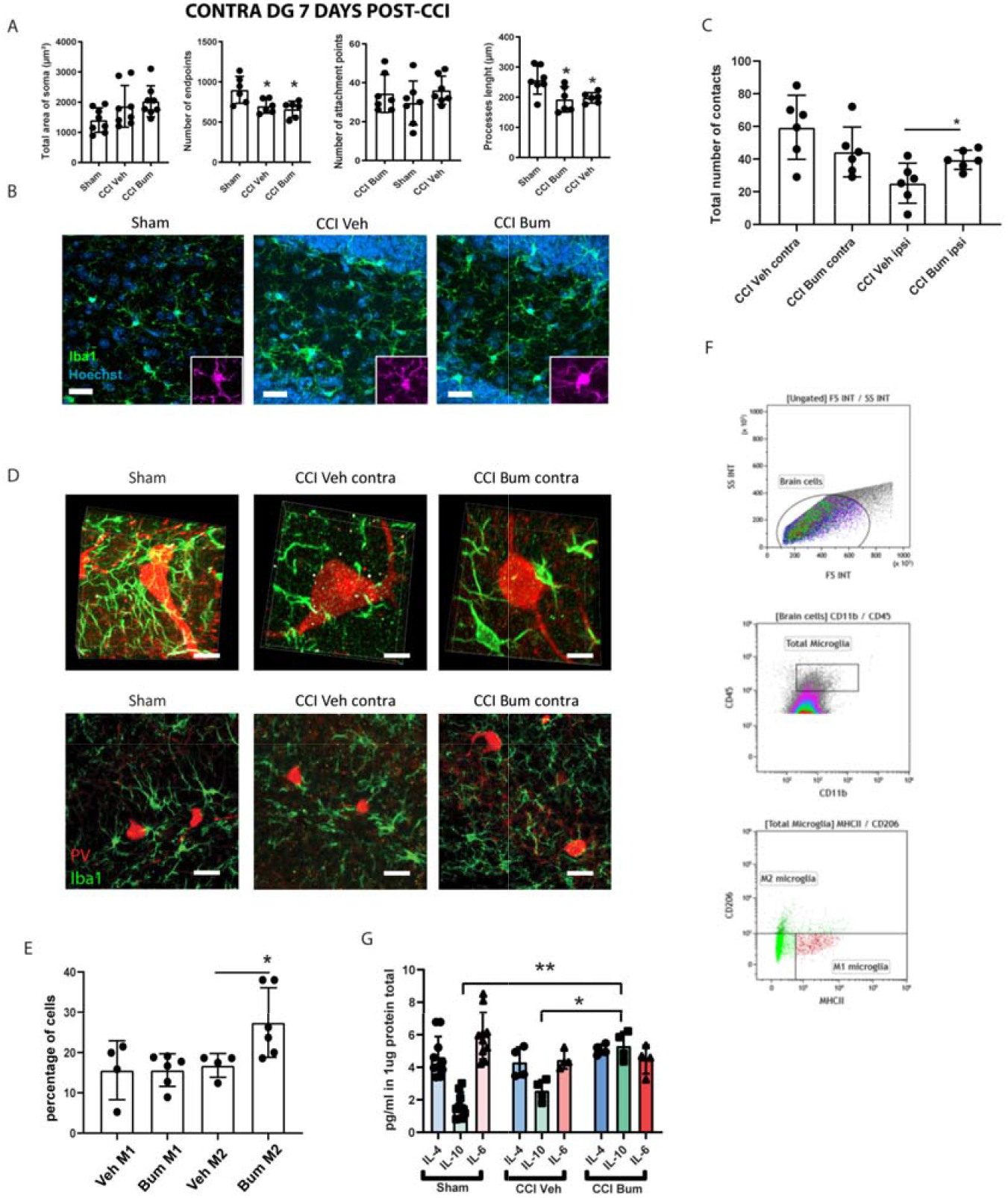
Microglia phenotypes and morphological changes after CCI and effect of bumetanide in the contralesional dentate gyrus at 7 days post-CCI. **(A**) Total area of Iba1+ cells soma, quantifications of Iba1 processes endpoints’, attachment points and length in the contralesional dentate gyrus 7 days post-CCI. n=5 animals, 2 slices per animal. **(B)** Iba1 immunostaining from sham, CCI-vehicle and CCI-bumetanide treated animals in the contralesional dentate gyrus. **(C)** Quantification of the number of contacts between microglia (Iba1 staining) and parvalbumin interneurons at 7 days post-CCI. n=5 animals per condition, 3 slices per animal. **(D)** Example of parvalbumin and Iba1 immunostaining, from sham, CCI-vehicle and CCI-bumetanide treated animals in the contralesional dentate gyrus, 3D representations (upper pictures), and stack (lower pictures). **(E)** Flow cytometry gating plots at 7 days post-CCI in contralesional side. **(F)** Analysis of microglia phenotypes by detection of MHCII^high^ (M1) and CD206^high^ (M2) markers by flow cytometry in the contralesional side at 7 days post-CCI. n=5-6 animals per condition. **(G)** Quantity of IL-4, IL-10 and IL-6 produced in the contralesional hippocampus. Phenotype changes were analyzed by Welch Anova followed by T-test. Morphological analysis was quantified using the ImageJ plugins Neurphology and SynapCountJ and analyzed using a t-test with Welch’s correction. Interleukin production was analyzed using a Kruskall-Wallis test for IL-10 *p < 0.05; **p < 0.01; ***p < 0.001.

Interestingly at 7dpCCI we found only slight morphological changes in microglia cells with no effect of bumetanide in both ipsi and contralateral side: decrease of the number of endpoints (Sham 900.7 ± 167.9 vs CCI Veh 696.5 ± 86.72 vs CCI Bum 659 ± 100.4) and of the process’ length (Sham 256.7 µm ± 46.75 vs CCI Veh 197.8 µm ± 17.17 vs CCI Bum 192.7 µm ± 42.21) (Fig 5A and B). Despite this, the number of contacts between microglial cells and PV interneurons still differed for the ipsilesional side (CCI Veh 25.20 ± 12.30 vs CCI Bum 39.56 ± 5.85) (Fig 5C and D). Surprisingly, at this stage we found that bumetanide promoted microglia polarization towards a M2 phenotype (Veh M2 16.79 % ± 2.957 vs Bum M2 27.44 % ± 8.635, Fig 5E and F). In line with these results, we also found a significant increase of IL-10 expression in the contralesional side of CCI animals treated with bumetanide, compared with control and vehicle-treated animals (CCI Bum 5.32 pg/mL ± 0.86 vs Sham 1.69 pg/Ml ±0.78 and CCI Veh 2.57 pg/mL ± 0.65) (Fig 5G).

These results suggest that bumetanide protects PV interneurons by increasing contacts to microglia during the first post-traumatic week as well as by regulation of microglia phenotype. This allows us to postulate that bumetanide treatment, in addition to preserving secondary neurogenesis, may also protect PV interneurons from death, through an anti-inflammatory mechanism mediated by microglia.

### Bumetanide regulates post-CCI microglia process motility

To confirm our previous results showing that bumetanide influences microglial cells activation, we imaged Cx3CR1^+/GFP^ mice, following laser-induced injury on the parietal cortex, that were treated or not with bumetanide. We were able to study the process motility of microglia *in vivo* by live bi-photonic microscopy recording during the first week post-injury (Fig 6B and E). One day post laser injury, we did not observe a difference in the length of microglial processes or surveillance state between vehicle and bumetanide treated animals (Fig 6C and D). However, 5 days post injury, we found that microglial processes are significantly longer in bumetanide treated animals, and that the process motility is enhanced (Bum 2.98 ± 1.88 vs Veh 0,48 ± 0.47, Fig 6F and G). These experiments confirmed again that bumetanide has an influence on microglia’s surveillance state in accordance with the increase of number of contacts between microglia and PV-interneurons.

**FIG 6.** Effect of bumetanide on microglia surveillance in acute and chronic phase post laser injury. **(A)** Schematic representation of the experimental timeline. **(B)** Temporal color-coded projection of in vivo microglia live imaging 1 day post laser injury. **(C)** Normalized average dendritic length 1day post laser injury. **(D)** Standard deviation of dendritic length, 1 day post laser injury. (**E-G)** Same as in **(B-D**) but 5 days post laser injury. Acquisitions were made with a 2-photon microscope (Femtonics) and analyzed using Fiji software and the ImageJ plugin Neurphology. All data was analyzed using an unpaired T-test. *p < 0.05.

### Bumetanide induces NKCC1 and BDNF expression in microglia

To investigate how bumetanide can influence microglia morphology and their trophic actions we used BV2 murine microglial cell line. 24 hours of treatment with bumetanide did not show significant changes in NKCC1 expression (normalized value on control: Ctrl 1 ± 0.17 vs Bum 0.82 ± 0.29, n= 12 wells per conditions, Fig. 7A). However, after 72 hours we detected a significant increase (Ctrl 1 ± 0.18 vs Bum 2.24 ± 0.99, Fig 7A). We then assessed the morphology of BV2 cells. We observed that 24h of bumetanide treatment induced a significant increase of the average size of cells (Bum 3161 µm^2^ ± 371.3 vs Ctrl 1993 µm^2^ ± 164.1, Fig 7B and D). At 72h this effect was not significant (Bum 3100 ± 414.1 vs Ctrl 1974 ± 231.5, Fig 7E and G). The effect of bumetanide on the number of processes was the opposite. While no effect was found at 24h (Bum 15.90 ± 4.73 vs Ctrl 22.08 ± 6.96 Fig 7B and C), bumetanide induced a significant decrease after 72h (Bum 14.59 ± 5.205 vs Ctrl 35.16 ± 10.84, Fig 7F and G). These results further indicates that bumetanide can regulate microglial morphology by modulating chloride co-transporter NKCC1 expression.

**FIG 7.**
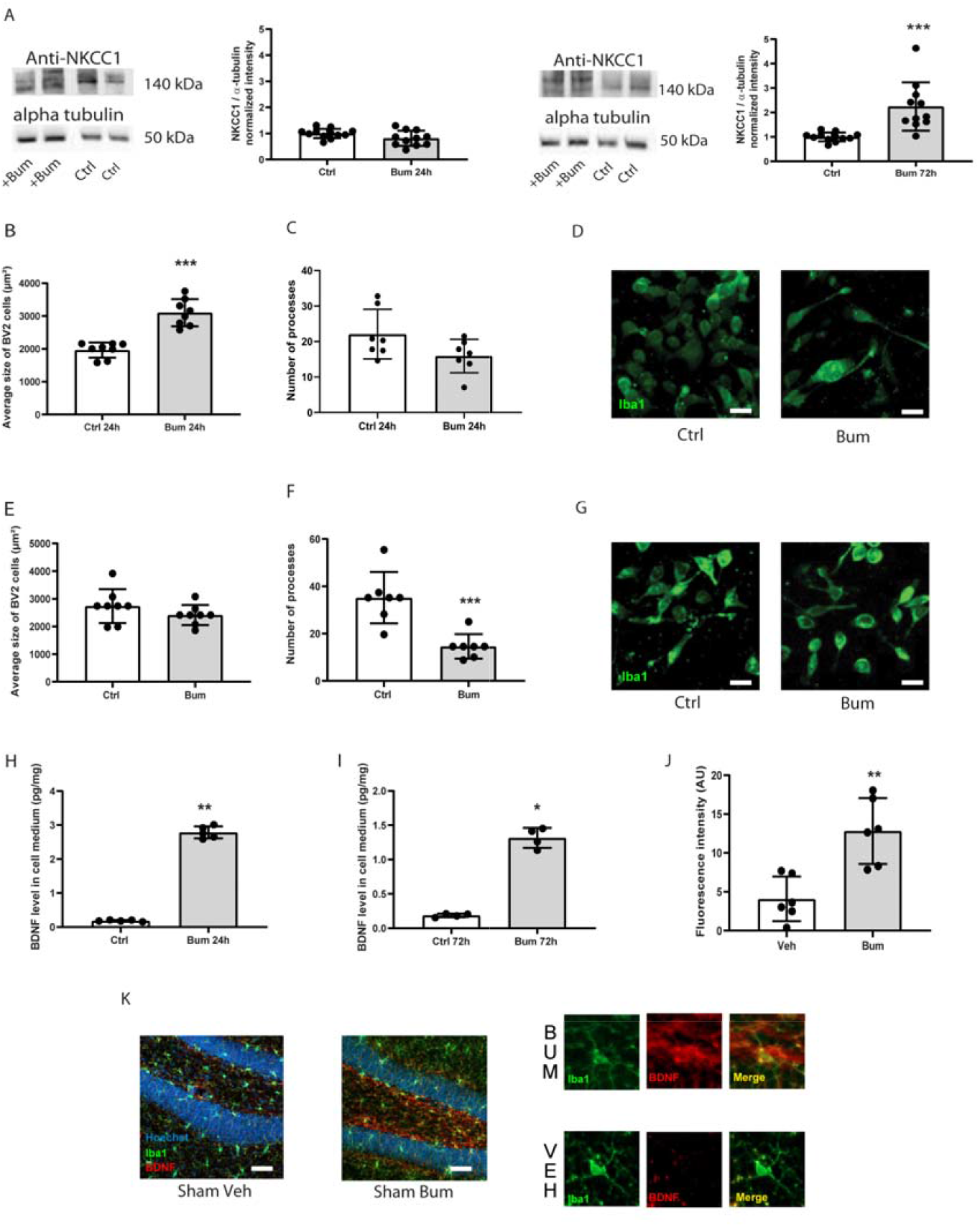
Microglia expresses NKCC1, its morphology and BDNF production is modified by LPS and bumetanide treatment *in vitro*. **(A)** The left panel represents NKCC1 protein expression normalized to the ubiquitous marker α-tubulin in BV2 cells after 24 hours of LPS, Bumetanide or LPS + Bumetanide treatment. On the right panel, NKCC1 normalized expression is shown after 72 hours of the same treatments. **(B)** Average size of BV2 soma in control condition and after 24 hours of bumetanide treatment. **(C)** Number of processes by BV2 cell in control condition and after 24 hours bumetanide treatment. **(D)** Iba1 immunostaining of BV2 cells cultured and fixed on labteck in control condition and after 24 hours bumetanide treatment. **(E)** Same as **(B)** after 72 hours of treatment. **(F)** Same as **(C)** after 72 hours of treatment. **(G)** Same as **(D)** after 72 hours of treatment. Measurement of BV2 BDNF levels after 24 **(H)** and 72 hours **(I)** of treatments. **(J)** Quantification of fluorescence intensity of BDNF staining in microglial cells. **(K)** Example of Iba1 and BDNF staining in dentate gyrus of Sham animals treated with bumetanide and non-treated. All sets of data were analyzed using Brown-Forsythe and Welch ANOVA tests followed by a Kruskal-Wallis post hoc test for NKCC1 expression, and BDNF levels analysis and Dunnett’s T3 multiple comparisons test for the BV2 morphology analysis. BDNF level of fluorescence were analyzed by a Mann-Whitney test. *p < 0.05; **p < 0.01; ***p < 0.001.

Microglia is a central source of BDNF which is an important neurotrophin controlling neuronal survival, and plasticity ^27,28^. Therefore, we measured the level of BDNF expression in BV2 cells after 24 and 72 hours of treatment. Surprisingly, 24 hours of bumetanide treatment led to a significant increase of BDNF levels (Ctrl 0.18 pg/mL ± 0.02 vs Bum 2.79 pg/mL ± 0.18, Fig 7H). This effect was attenuated but still present after 72 hours of treatment (Ctrl 0.19 pg/mL ± 0.03 vs Bum 0.31 ± 0.14, Fig 7I). Furthermore, we quantified the immuno-like intensity of BDNF in sham vehicle and sham bumetanide treated conditions and we found a consistent increase in BDNF expression within iba1 positive microglia (Sham Veh 4.093 ± 2.87 vs Sham Bum 12.82 ± 4.27, Fig 7J and K). These results indicate that bumetanide may have direct effect on microglia, and notably induce its BDNF production that may play an important role in the survival of PV interneurons.

## Discussion

How cognitive deficits observed in experimental models and clinical TBI are formed following an initial concussion is not well understood ^3^. In this study, we were interested in how chloride homeostasis is involved in the mechanism that could potentially target changes in working memory ^29^. Theta rhythm is a potential electrophysiological biomarker of altered neuronal activity ^30,31^ and is believed to synchronize activity both within local networks and between distal cortical regions involved in cognitive processing ^32^. In both humans and rats, the power of theta rhythm increases during the acquisition phase of spatial learning and object recognition. Furthermore, high-power theta oscillations are predictive of success on spatial cognitive tasks and, in particular, spatial learning ^33^. Treatment-or injury-induced inhibition of theta oscillations results in cognitive dysfunction ^22^. TBI in rodents decrease the theta/gamma ratio and correlates with cognitive deficits ^34,35^. The behavioral results in the present study fit well to this hypothesis as we found a positive correlation between decreased theta-band power and poor performance in spatial cognitive test. Interestingly, we found that an early transient treatment with the chloride cellular uptake inhibitor bumetanide was effective to counteract changes of both behavior and theta oscillations.

Several studies link cognitive processes with normal adult neurogenesis in the hippocampus. This corresponds to the generation of new functional neurons from adult neural stem cells through the amplification of intermediate progenitors and neuroblasts. TBI can induce significant changes in the proliferation and maturation of granular cells ^36,37^. In the present study we found a significant increase in the number of Nestin positive cells in both ipsi and contralesional dentate gyrus of mice 7 days following CCI. Nestin is a radial glia cell marker (RGL), and represent a pool of quiescent cells that divide only occasionally ^16^. The division of these RGLs leads to the generation of intermediate progenitor cells (IPCs) or type II cells that will undergo a limited number of rapid divisions before leaving the cell cycle and entering the neuronal differentiation pathway ^38^. Type II cells are divided into two subtypes: i) type 2a cells that are positive for Nestin and negative for double-cortin (DCX), an immature neuronal marker, and ii) type 2b cells, positive for Nestin and DCX ^39^. IPCs then give rise to neuroblasts (type 3 cells), which express DCX ^40^. The increased number of Nestin positive cells after TBI might intuitively suggest that there will be an increase in the number of newborn neurons. This is however not the case since we found a significant decrease of DCX positive cells, in both the contra- and ipsilesional side of the DG. These results indicate that TBI leads to an increase of the quiescent cell pool and a resulting decrease in number of immature granule cells consistent with previous findings^6^. Furthermore, treatment with bumetanide during the first week post TBI significantly reduced the CCI induced decrease in Nestin positive cells as well as increased the number of DCX positive cells. These results suggest that the effects of bumetanide on cognitive performance could be related to positive effects on post-traumatic granule cells production. In addition, they may suggest a common upstream mechanism for the action of bumetanide.

GABA, especially released by PV interneurons ^7,41^, appears to play a central role in regulating the activity of RGLs by inhibiting their proliferation. Thus, post-traumatic changes in granule cells could be related to abnormal interneuron function. Indeed, we found loss of parvalbumin interneurons in the granular layer of the hippocampus, as early as 3 days post-CCI in both the ipsi and contralesional side that was sustained at 7 days post CCI and one month after trauma^6^.This loss of parvalbumin interneurons may also contribute to theta-band changes after TBI ^42^.

The increase of Nestin positive cells that we detected after TBI can thus be related to the loss of PV interneurons that results in less ambient GABA and downstream unlock of the pool of quiescent Nestin positive cells. As tonically GABAergic activation by ambient GABA is considered to be the initial inputs of these type 2 progenitors and is thought to play a role in their maturation it is plausible that the decrease of newborn neurons can also be explained by changes in PV derived GABA ^43,44^. In accord with this assumption, treatment with bumetanide significantly counteracted the loss of PV interneurons.

Neuroinflammation is observed in both the acute and chronic stages of TBI^12^. It appears to be responsible for both, detrimental and beneficial effects, by contributing to the primary injury and secondary damage, but also facilitating tissue repair^45^. Both astrocytes and microglia are very important players in the mechanisms of brain inflammation and in addition display one of the highest expression levels of NKCC1 in the brain. Thus is plausible that the post-traumatic effects produced by bumetanide could be related to an influence on the dynamics of inflammatory processes^45,46^. In support of this assumption, treatment with a derivative of tetracycline widely used to reduce inflammation, Minocycline^47^, resulted in similar effects than those found with Bumetanide e.g. Minocycline also led to an increase in the number of newborn neurons and rescue of PV interneurons.

Following this idea, we first studied the role of NKCC1 mediated chloride uptake in astrocytes using conditional transgenic animals. Despite bumetanide having significant effects on the morphology of GFAP positive cells in both contra and ipsilateral side in sham animals, the ablation of NKCC1 in these cells did not induce significant rescue effects on the survival of PV positive interneurons. This indicates that astrocytes are not significantly involved in the bumetanide-induced increment in interneuron survival.

As astrocytes were not significantly involved in the effect of bumetanide we then chose to focus on microglia, the main resident immune cells of the brain. In a first step, we were keen to know if neuroinflammation through microglia could have an effect on the PV interneuronal loss. To this end, we found that bumetanide treatment leads to more contact between microglia and PV interneurons. This is consistent with previous results showing pro-survival signaling mediated by microglia contact on interneurons ^48^. These results imply that the mechanism by which bumetanide rescues PV interneurons can be related to additional effects on microglia morphology. Our experiments on microglia morphology confirmed this and showed that the treatment leads to a significant increase of microglial processes and ramifications as well as an increase of the cell soma 3 days post-CCI. Moreover, the bumetanide treatment significantly changed microglia processes motility *in vivo*.

In addition to morphological changes, bumetanide treatment also induced phenotypic changes in microglia. We showed that bumetanide treatment leads to a faster activation of microglial cells into a pro-inflammatory phenotype M1 at early stages, which is required for cell debris removal and phagocytosis of apoptotic cells. At 7 days post-CCI, microglial cells of bumetanide treated animals acquired the M2 phenotype. Microglia in control animals acquired the M1 phenotype slightly later but above all, this pro-inflammatory phenotype persists more than 7 days after trauma, resulting in a deleterious inflammatory state. The fact that bumetanide increased the concentration of interleukin 6, 3 days post-CCI is interesting. A recent study, using the same trauma model, showed that IL-6 production by hippocampal granule cells in the presence of renewed microglial cells would induce neuronal survival and would notably allow adult neurogenesis to proceed normally ^49^. The presence of IL-6 would also prevent the behavioral deficits ^49^. One-week post-CCI, we detected higher levels of interleukin 10 in bumetanide-treated animals which goes along with the M2 phenotype switch that we detected. Expression of IL-10 can promote neuronal and glial cell survival and dampen inflammatory responses via a number of signaling pathways ^50^. To further investigate the effect of bumetanide on microglia, we used the BV2 cell line *in vitro* and observed morphological changes after bumetanide treatment. Importantly, we found a significant increase of the neurotrophic factor BDNF. Increased BDNF release in combination with increase contact could mediate an increased survival of PV interneurons.

This study shows for the first time that a treatment targeting the chloride co-transporter NKCC1 can regulate the activation kinetics of microglia and provides mechanistic link for the positive effect of bumetanide on post-traumatic cognitive decline. These results also open an additional path to better understand the pathophysiological mechanisms in the post-traumatic brain through cellular chloride uptake and help to understand the factors involved in the development of long-term sequelae following TBI.

## Acknowledgments

This work was supported by public Aix-Marseille Université (AMU); Eranet Neuron III program through the Acrobat, ANR GABGANG project and Academy of Finland project nr 341361 to CR; and by the “fondation des gueules cassées” through MT. We thank Prof Eero Castren and Yves-Alain Barde for providing the BDNF antibody.

## Author Contributions

MT performed research (TBI model, IHC, biochemistry, morphological analysis, cell culture, surgery)/ analyzed data / wrote the manuscript. MSG performed the live imaging experiment, LT performed the flow cytometry experiment, KC analyzed the flow cytometry data, FM helped planning and designing the experiments on hGFAP NKCC1 flox mice, EG helped injecting mice and performing the ELISA experiment, FG provided the Nestin-GFP mice, CH provided the NKCC1 flox mice, JL revised the manuscript, CP designed research / performed the behavioral analysis / revised the manuscript, CR designed research / revised the manuscript

